# The regulatory landscape of the human HPF1- and ARH3-dependent ADP-ribosylome

**DOI:** 10.1101/2021.01.26.428255

**Authors:** Ivo A. Hendriks, Sara C. Buch-Larsen, Evgeniia Prokhorova, Alexandra K.L.F.S. Rebak, Ivan Ahel, Michael L. Nielsen

**Author notes:** Equal contribution (Ivo A. Hendriks, Sara C. Buch-Larsen).

## Abstract

Despite the involvement of Poly(ADP-ribose) polymerase-1 (PARP1) in many important biological pathways, the target residues of PARP1-mediated ADP-ribosylation remain ambiguous. To explicate the ADP-ribosylation regulome, we analyzed human cells depleted for key regulators of PARP1 activity, histone PARylation factor 1 (HPF1) and ADP-ribosylhydrolase 3 (ARH3). Using quantitative proteomics, we quantified 1,596 ADPr sites, displaying a thousand-fold regulation across investigated knockout cells. We find that HPF1 and ARH3 inversely and homogenously regulate the serine ADP-ribosylome on a proteome-wide scale with consistent adherence to lysine-serine (KS)-motifs suggesting targeting is independent of HPF1 and ARH3. Our data reveal that ADPr globally exists as mono-ADP-ribosylation, and we detail a remarkable degree of histone co-modification with ADPr and other post-translational modifications. Strikingly, no HPF1-dependent target residue switch from serine to glutamate/aspartate was detectable in cells, which challenges current dogma related to PARP1 target residues. Collectively, we elucidate hitherto unappreciated processes related to cellular PARP1 activity.

## Introduction

ADP-ribosylation (ADPr) is catalyzed by poly-ADPr-polymerases (PARPs), also known as ADP-ribosyltransferases (ARTs). ADPr refers to the process where an ADP-ribose moiety is transferred from NAD+ to the amino acid side-chains of target proteins (Schreiber et al., 2006). Although discovered more than 50 years ago (Chambon et al., 1963), neither the biological functions of PARP1-catalyzed ADPr nor its contributions to cell biology are fully understood. ADPr imposes structural constraints and introduces negative charges to acceptor proteins, thereby altering protein localization, protein function, and interactions with other proteins and DNA (Ray Chaudhuri and Nussenzweig, 2017). Due to this, PARP1 has emerged as a master regulator in the DNA damage response through its crux position in the base excision repair pathway (Schreiber et al., 2006). Given the importance of ADPr in the DNA repair process, inhibitors of catalytic PARP activity (i.e. PARP inhibitors) are widely used in the clinic to combat various cancers. Hence, new insight into the enzymatic catalysis of ADPr advances our understanding of PARP biology, and refines current knowledge related to the mechanisms surrounding sensitivity and resistance to PARP inhibitors in cancer (Lord and Ashworth, 2012, 2013).

Despite the biomedical importance of PARP1-mediated ADPr, and inhibition thereof, the molecular details surrounding which amino acid residues are PARP1 targets has remained an analytical conundrum (Crawford et al., 2018; Leung, 2017). PARP1-catalyzed ADPr has historically been reported to target the side-chains of glutamate and aspartate, both as PARP1 auto-ADPr (Tao et al., 2009), and on downstream protein substrates (Daniels et al., 2014; Ogata et al., 1980; Zhang et al., 2013). Recently, serine residues were reported as the major target for PARP1-mediated ADPr in human cells during DNA damage (Hendriks et al., 2019; Larsen et al., 2018; Leidecker et al., 2016; Palazzo et al., 2018), under physiological conditions (Buch-Larsen et al., 2020), and in the context of PARP1 auto-ADPr (Bonfiglio et al., 2017; Larsen et al., 2018). While the mechanistic details surrounding the catalytic preferences of PARP1 remain perplexing, observations that PARP1/2 activity is regulated by Histone PARylation factor (HPF1) (Gibbs-Seymour et al., 2016) has provided some biochemical and structural understanding to this ambiguity (Bilokapic et al., 2020; Suskiewicz et al., 2020b). *In vitro* experiments support that availability of HPF1 changes the catalytic preference of PARP1 from glutamate and aspartate to serine residues (Blessing and Ladurner, 2020; Bonfiglio et al., 2017; Suskiewicz et al.). However, an unbiased investigation of an HPF1-dependent switch in target residues from serine to glutamate and aspartate in cells is currently lacking.

The catalytic activity of HPF1-PARP1/2 is counteracted by poly-ADP-ribose glycohydrolase (PARG) (Slade et al., 2011), and ADP-ribosylhydrolase 3 (ARH3) (Abplanalp et al., 2017; Fontana et al., 2017), providing an additional layer of complexity to the cellular ADPr dynamics. This leads to high turnover of poly-ADP-ribosylation (PAR) and certain PARP1-catalyzed modifications existing as mono-ADPr (MAR) events in human cells (Bonfiglio et al., 2020; Fontana et al., 2017). With PARP1 described as a poly-ADPr (PAR) polymerase (Kraus, 2015; Schreiber et al., 2006), the notion that PARP1 substrates primarily exist as MARylated entails fundamental biological insights and epitomizes the relevance of exploring the modularity of ADPr under *in vivo* conditions. However, at present, only a limited number of *in vivo* substrates have been described as MARylated. Hence, comprehensive and unbiased analyses are required to assess whether MARylation is a global phenomenon, considering that PARP1 is able to target hundreds of protein substrates (Hendriks et al., 2019).

To alleviate these knowledge gaps, while concomitantly providing a valuable resource to the community on the enzymatic modularity of the ADP-ribosylome, we employed an Af1521-based proteomic approach (Martello et al., 2016) to compare ADPr acceptor sites and protein substrates across human cells genetically depleted for HPF1 or ARH3 (Fontana et al., 2017; Gibbs-Seymour et al., 2016). Our results support that HPF1 and ARH3 are global regulators of serine ADPr, and we outline that the absence of HPF1 does not result in increased ADPr on glutamate and aspartate residues. Collectively, our data summarizes the HPF1- and ARH3-mediated ADPr regulation at a proteome-wide scale, and provides new insights into the cellular distribution of PARP1-catalyzed ADPr target residues.

## RESULTS

### Mono-ADP-ribosylation is predominant in human cells

With the Af1521 macrodomain able to bind both mono- and poly-ADP-ribose (Karras et al., 2005; Rosenthal et al., 2013), we reasoned that moderations to our proteomics enrichment strategy would allow for assessing the global extent of MAR in human cells (Fig. 1A) (Larsen et al., 2018). While macrodomains are able to hydrolyze ADP-ribose moieties (Rack et al., 2016), we previously demonstrated that the Af1521 macrodomain does not exert hydrolase activity (Jungmichel et al., 2013). Nonetheless, we wanted to ensure that our MS-based proteomics strategy using Af1521 enrichment preserves all ADPr and is unbiased with regard to identification of amino acid acceptor sites. To this end, we compared Af1521 to two independent antibodies (from Cell Signaling Technology) raised against poly/mono-ADP-ribose, as antibodies should not possess hydrolase activity and thus allow unbiased enrichment of ADP-ribosylation for proteomics analysis (Fig. 1A). PARG is integral to our established protocol (Hendriks et al., 2019), but might exhibit off-target activity. To control for this, we evaluated purification of ADPr without addition of PARG, or with addition of ARH3, which should remove all serine ADPr (Abplanalp et al., 2017) (Fig. 1A). We used our contemporary method to purify ADPr from quadruplicate HeLa cell cultures exposed to oxidative stress, and analyzed the ADPr peptides by high-resolution mass spectrometry, relying on the non-ergodic fragmentation propensity of electron-transfer dissociation (ETD) (Coon et al., 2005) to allow for faithful localization of the PTM to the correct amino acid (Larsen et al., 2018).

**Figure 1.**
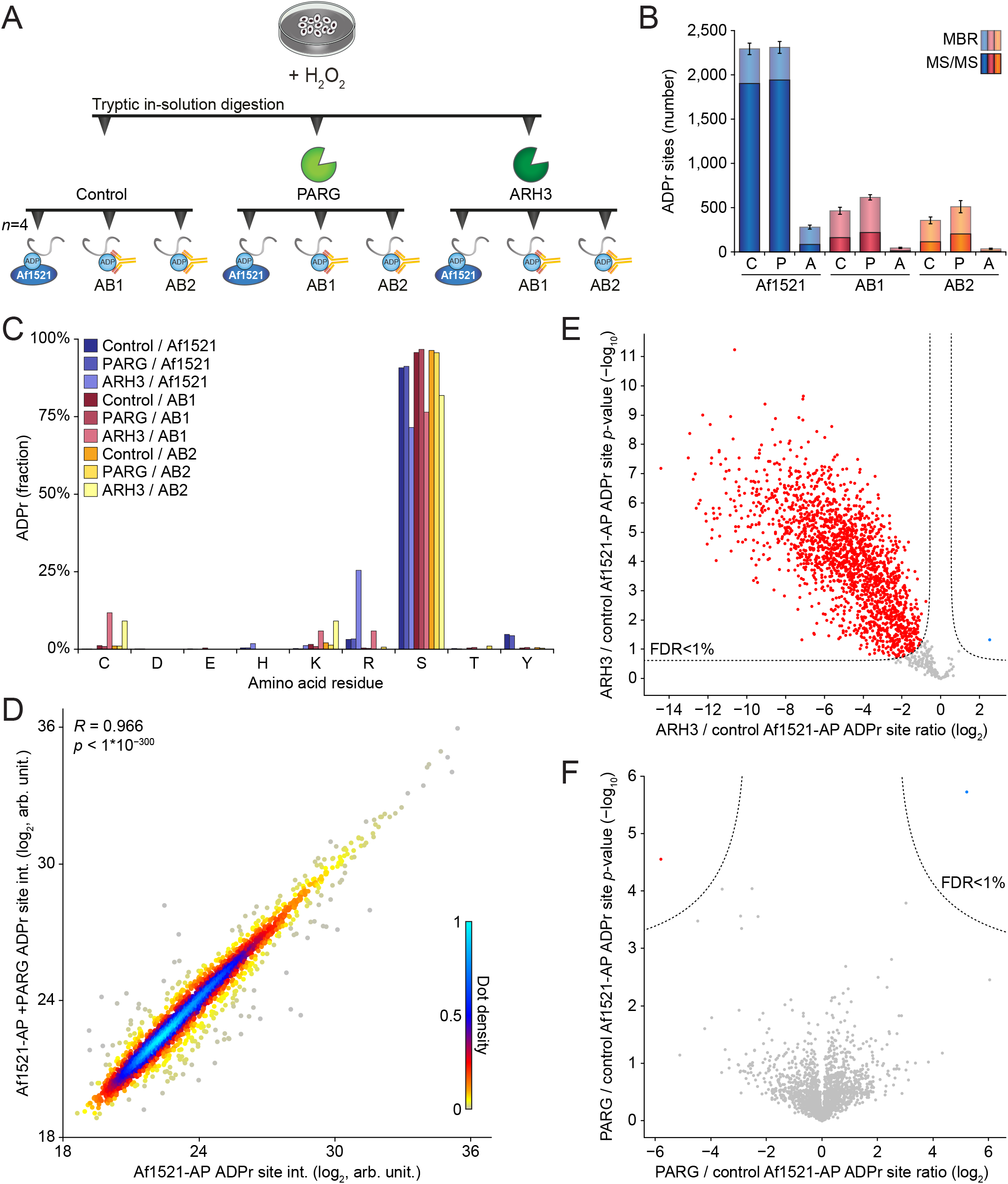
Evaluation of different erasers and enrichers for the purification of ADPr. (A) Overview of the experimental design. HeLa cells were cultured, H_2_O_2_-treated at 1 mM for 10 min, lysed, and either mock treated, PARG-treated, or ARH3-treated. Quadruplicate (*n*=4) ADPr peptide purifications were performed using either Af1521 macrodomain, E6F6A antibody (AB1), or D9P7Z antibody (AB2). ADPr peptides were analyzed using high-resolution MS. (B) Overview of the number of ADPr sites identified and localized (>90% probability). Error bars represent SD, *n*=4 purification replicates. “C”; control, “P”; PARG, “A”; ARH3, “MBR”; matching between runs. (C) Visualization of the abundance fraction of ADPr as distributed across different amino acid residue types. (D) Scatter plot analysis demonstrating correlation between mock treated and PARG-treated ADPr sites. “R” indicates Pearson correlation, p-value determined via linear regression t-test. (E) Volcano plot analysis comparing ARH3-treated versus mock treated Af1521-enriched ADPr sites. Red and blue dots indicate significantly down- and upregulated sites, respectively, with permutation-based FDR-corrected p-value <1%. (F) As **E**, but comparing PARG-treated versus mock treated Af1521-enriched ADPr sites.

In total, we identified and localized 2,758 ADPr sites (Table S1), of which the majority was identified via the Af1521 purification (Figs. 1B and S1A). Both antibodies allowed direct identification of ∼200 ADPr sites, and up to ∼500 by matching MS1-level evidence from Af1521 runs. With all three purification methods, ADPr was primarily detected on serine residues (Fig. 1C), supporting that the Af1521 methodology is unbiased. In line with expectations, *in vitro* ARH3 treatment of peptides prior to ADPr purification greatly reduced the number of identified ADPr sites (Fig. 1B), with non-serine ADPr becoming relatively more abundant after *in vitro* ARH3 treatment (Fig. 1C). Strikingly, omitting *in vitro* PARG treatment did not notably alter the number of ADPr sites detected (Fig. 1B), and we observed a strong correlation between ADPr site abundances from untreated and PARG-treated samples (Figs. 1D and S1B). Furthermore, whereas we observed a significant decline in the majority of all ADPr sites upon *in vitro* ARH3 treatment (Fig. 1E), this was not the case after PARG treatment (Fig. 1F), indicating that *in vitro* PARG treatment was redundant for reduction of ADPr polymer length. Collectively, we demonstrate that our Af1521 methodology is unbiased, and our results support that mono-ADPr is a global phenomenon in cells.

### HPF1 and ARH3 globally regulate a homogenous ADP-ribosylome

The important and contrasting roles of HPF1 and ARH3 in serine ADPr homeostasis were previously elucidated (Abplanalp et al., 2017; Bonfiglio et al., 2017; Fontana et al., 2017; Gibbs-Seymour et al., 2016; Suskiewicz et al., 2020b). However, insight into the systemic effect of these enzymes on global and site-specific ADP-ribosylation is currently lacking. To this end, we cultured either wild-type (control) U2OS cells or U2OS lacking HPF1 or ARH3 in quadruplicate. To capture the dependency of HPF1 and ARH3 across various biological processes, we analyzed cells both under oxidative stress known to induce ADP-ribosylation (Luo and Kraus, 2012), and untreated steady-state conditions (Fig. 2A). We validated the absence of HPF1 or ARH3 via immunoblot analysis (Fig. 2B), and confirmed strong induction of ADPr in response to H_2_O_2_ treatment and ARH3 depletion (Fig. 2C). Subsequently, we purified ADPr sites from all replicate cultures using our established and unbiased Af1521 methodology, and analyzed them via mass spectrometry. Overall, we identified 1,596 ADPr sites (Table S2), corresponding to 799 ADPr target proteins (Table S3). On average, the largest numbers of ADPr sites were observed in ARH3 KO cells (Fig. 2D), even in the absence of H_2_O_2_ treatment), which closely resembled immunoblot observations (Fig. 2C). In terms of ADPr abundance, HPF1 KO nearly abolished detectable ADPr, whereas ARH3 KO resulted in dramatic ADPr accumulation, with ∼100- and ∼1,000-fold increases compared to control and HPF1 KO cells, respectively (Fig. 2E). We observed a high degree of replicate reproducibility (Figs. 2F-G), and a strong tendency for untreated control cells to resemble HPF1 KO cells, with H_2_O_2_-treated control cells resembling ARH3 KO cells. Indeed, H_2_O_2_ induction of ADPr was predominantly observed in wild-type cells (Figs. 2D-F and 2H-I), whereas the KO cell lines responded markedly less to H_2_O_2_ treatment. A considerable number of ADPr sites induced in response to H_2_O_2_ treatment in control cells were detectable in untreated ARH3 KO cells (Figs. 2F), and the majority of control ADPr sites were also detectible in the ARH3 KO cells (Figs. 2H-I). Homogeneity between untreated ARH3 KO cells and H_2_O_2_-treated control cells was supported by Pearson correlation analysis of identified ADPr modification sites (Fig. S2A), and term enrichment analysis on ADPr target proteins detected under both conditions revealed many canonical features associated with ADP-ribosylation (Fig. S2B), such as nuclear and chromosomal localization, and modification of proteins involved in RNA metabolism and DNA repair. Overall, we find that ARH3 KO results in accumulation of vast amounts of ADPr, while remaining functionally homogenous with the majority of ADPr usually observed in response to oxidative stress.

**Figure 2.**
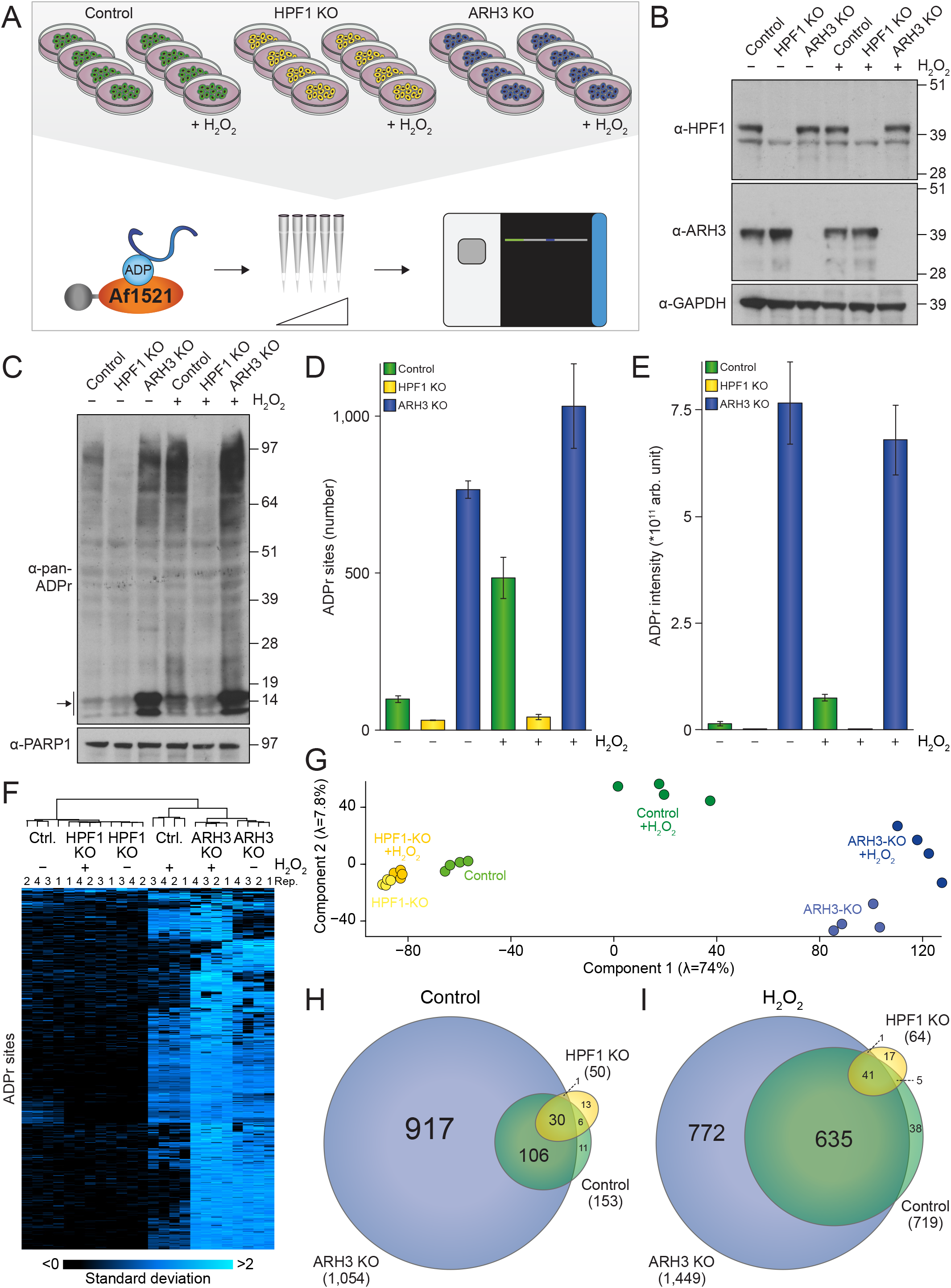
The ADP-ribosylome in HPF1 and ARH3 knockout cells. (A) Overview of the experimental design. U2OS cells, either wild-type (control), HPF1 knockout (KO), or ARH3 KO, were cultured in quadruplicate (*n*=4), and either mock- or H_2_O_2_-treated at 1 mM for 10 min. ADPr sites were enriched using the Af1521 methodology, fractionated, and analyzed using high-resolution MS. *n*=4 cell culture replicates. (B) Immunoblot analysis validating the knockout of HPF1 and ARH3. (C) Immunoblot analysis highlighting the ADPr equilibrium in control, HPF1 KO, and ARH3 KO cells, +/− H_2_O_2_ treatment at 1 mM for 10 min. The arrow indicates histone ADPr. (D) Overview of the number of identified and localized ADPr sites. *n*=4 cell culture replicates, error bars represent SD. (E) As **D**, showing ADPr abundance. (F) Hierarchical clustering analysis of z-scored ADPr site abundances, visualizing relative presence of ADPr sites across the experimental conditions. (G) Principle component analysis indicating the highest degree of variance between sample conditions. (H) Scaled Venn diagram depicting overlap between ADPr sites in untreated cells. (I) As **H**, but for H_2_O_2_-treated cells.

### Absence of HPF1 does not cause redirection of ADPr to other residue types

We have previously demonstrated that serine residues are the primary target of ADPr in cultured cells (Larsen et al., 2018), and the predominance of lysine-directed serine ADPr in the form of the lysine-serine (KS) motif (Hendriks et al., 2019; Leidecker et al., 2016). We scrutinized the prevalence of these phenomena in the context of HPF1 and ARH3 KO, and found that serine ADPr was overall predominant (Fig. 3A), with an increase to >99% abundance in ARH3 KO cells or in response to H_2_O_2_ in control cells, while even in HPF1 KO cells serine ADPr narrowly stayed in the lead. The large majority of serine ADPr resided within KS motifs (Fig. 3B), even in the absence of HPF1, suggesting that the preferential targeting of ADPr to the KS motif does not rely solely on HPF1. In terms of amino acid distribution, the number of serine ADPr sites was by far the greatest (Fig. 3C), corresponding to abundance-based observations (Fig. 3A). Notably, in the absence of HPF1, where the total ADPr signal is greatly reduced (Fig. 2E), the remaining ADPr was observed to also target histidine (∼24%), arginine (∼11%), and cysteine (∼8%) residues, with only weak evidence supporting trace ADPr targeting aspartic and glutamic acid residues (Fig. 3C). Substantiating these observations, treatment of cell lysates with hydroxylamine did not change the distribution of ADPr (Fig. S2C), furthering that aspartic and glutamic acid residues are not highly represented in any of the evaluated experimental conditions. Directly comparing the sequence context of ADPr between HPF1 and ARH3 KO cells revealed no significant changes in amino acid distribution directly surrounding the ADPr sites (Fig. 3D), and highlighted a significant shift in modification away from serine residues and primarily towards histidine, cysteine, and arginine residues. Notably, this shift in preference away from serine was relative, as the absolute number of non-serine ADPr sites did not significantly change (Table S2).

**Figure 3.**
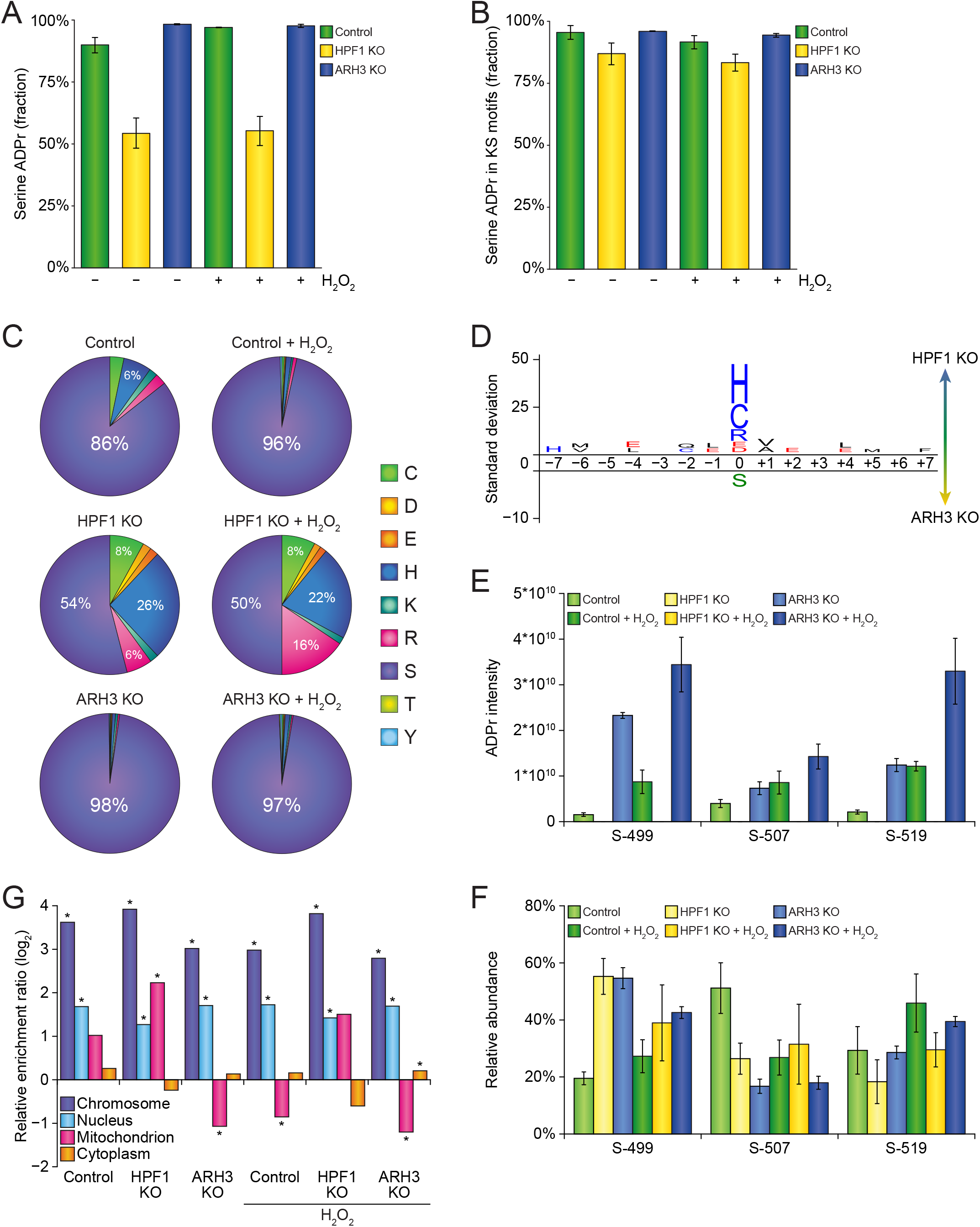
Site-specific properties of ADPr in HPF1 and ARH3 knockout cells. (A) Overview of the abundance fraction of ADPr modifying serine residues. *n*=4 cell culture replicates, error bars represent SD. (B) As **A**, but visualizing the fraction of serine ADPr in (lysine-serine) KS motifs. (C) Pie-chart analysis showing the distribution of ADPr sites across different amino acid residue types. (D) IceLogo analysis visualizing relative preference for ADPr to be targeted to different amino acid residue types. Amino acid residues displayed above the line were enriched for HPF1 KO, and those displayed below were enriched for ARH3 KO. All displayed amino acids indicate significant changes as determined by two-tailed Student’s t-testing, *n*= 50 and *n*=1,054 HPF1 KO and ARH3 KO ADP-ribosylation sites, respectively, *p*<0.05. (E) PARP1 auto-modification analysis, showing absolute modification abundance. *n*=4 cell culture replicates, error bars represent SEM. (F) As **E**, but showing relative modification abundance. (G) Term enrichment analysis visualizing the Gene Ontology subcellular localization of ADPr target proteins across experimental conditions. Significance was determined via Fisher Exact testing with Benjamini-Hochberg correction for multiple hypotheses testing, \**p*<0.05.

PARP1 is mainly modified on S-499, S-507, and S-519, residing in the auto-modification domain (Bonfiglio et al., 2017; Larsen et al., 2018; Leidecker et al., 2016). Here, we observed that the abundance of PARP1 auto-ADPr (Fig. 3E) followed the global trend for total ADPr with regard to ARH3 KO and H_2_O_2_ treatment (Fig. 2E). Under all conditions, including HPF1 KO, we noted predominant targeting of ADPr to the three primary serine residues (Fig. 3F). In terms of subcellular localization of proteins modified by ADPr, a strong enrichment was observed for chromosomal and nuclear localization across all experimental conditions (Fig. 3G). We did not note a particular preference for cytoplasmic ADPr target proteins; however, in the absence of HPF1 and in untreated control cells, we noted a relative preference for ADP-ribosylation of mitochondrial proteins (Fig. 3G), corroborating our previous observations of histidine ADPr targeting the mitochondria (Hendriks et al., 2019). Taken together, the absence of HPF1 resulted only in reduction of serine ADPr, rendering non-serine ADPr relatively but not absolutely more prominent.

### Histone variants primarily contain serine ADPr

The predisposition of histones to be ADP-ribosylated has been well-characterized (Burzio et al., 1979; Jump et al., 1979; Messner and Hottiger, 2011). In ARH3 KO cells, we found a striking prevalence of histone ADPr (Fig. 4A), whereas the relative fraction of histone ADPr was not notably altered in control and HPF1 KO cells, or in response to H_2_O_2_ treatment. Investigation of modification across all histone variants unveiled the highest degree of modification on H3 and H2B (Figs. 4B and S3A), in accordance with previous *in vivo* observations (Bartlett et al., 2018; Boulikas, 1988; Rakhimova et al., 2017). On all histone variants, we noted that a single ADPr site accounted for the vast majority of total modification (Fig. 4B-C). Intriguingly, owing to the elevated levels of histone ADPr in the absence of ARH3, we found that Histone H1 constitutes the third-most ADPr-modified variant. Following this, we aligned the sequences of all Histone H1 isoforms, and mapped all identified ADPr sites to the alignment (Fig. 4D). Our deep MS analysis facilitated direct identification of ADPr on Histones H1.0, H1.1, H1.2, H1.4, H1.6 (H1t), and H1.10 (H1x), with non-unique peptides also mapping to H1.3 and H1.5 (Fig. 4D). Immunoblot analysis was performed on GFP-tagged H1 isoforms, validating ADPr on H1.0, H1.1, H1.2, H1.3, and H1.4 (Figs. 4E and S3B). With H1.5 not detected via unique peptides in the MS and not validated via IB, we suggest that H1.5 is not modified by ADPr. Whereas the absence of HPF1 abolished ADPr on Histone H1 (Fig. 4E), absence of ARH3 conversely increased histone H1 ADPr levels (Fig. S4A). Overall, in terms of modification abundance, H1.4 was the primary target of ADPr, followed by H1.0 and H1.2 (Fig. 4F). Site-directed mutagenesis of four main serine ADPr sites (>98% abundance as detected by MS) in Histone H1.2 eliminated detectable ADPr (Fig. S4B), confirming our MS data and collectively highlighting that virtually all ADPr on histones occurs on serine residues.

**Figure 4.**
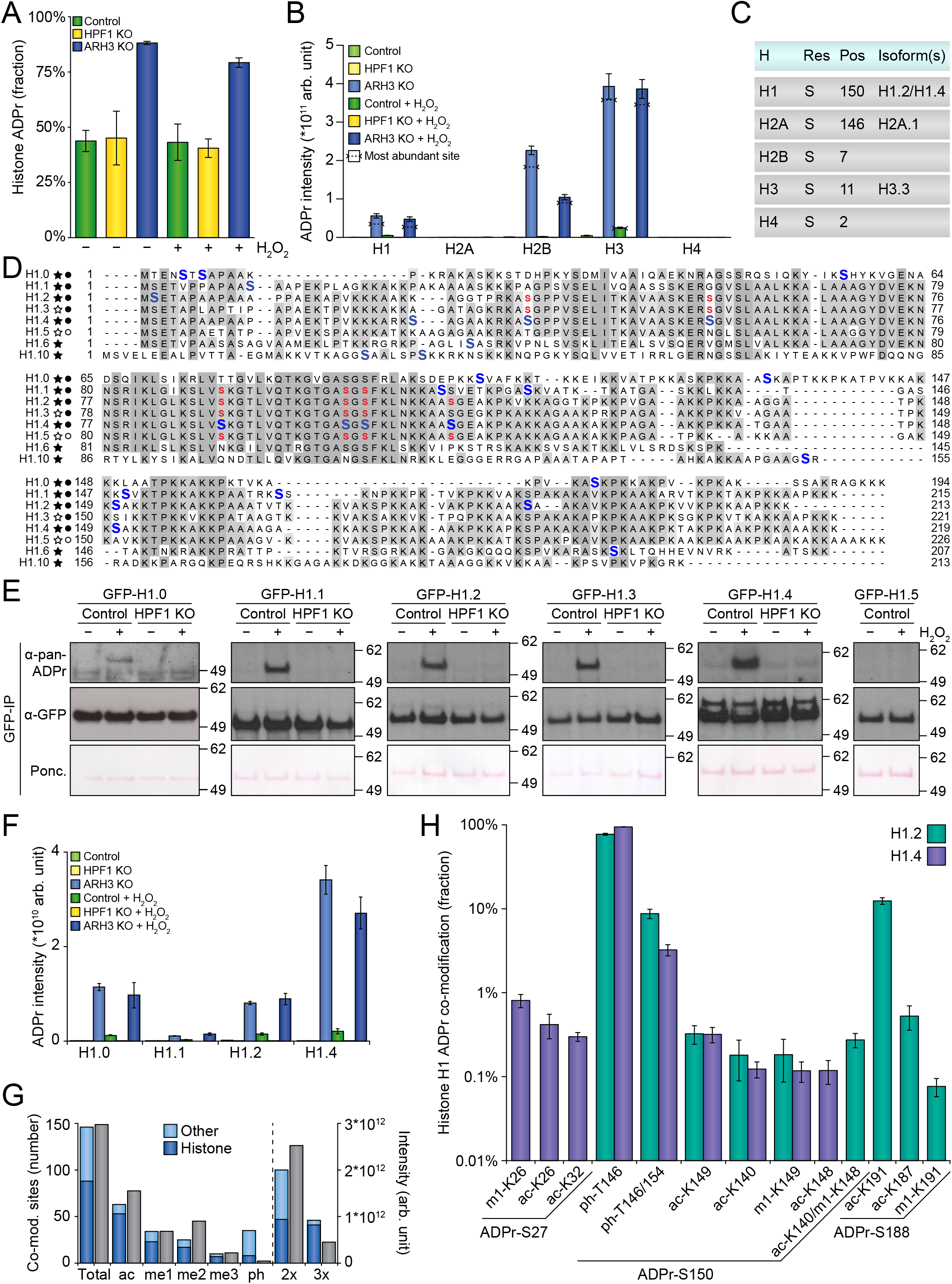
A comprehensive analysis of the histone ADPr landscape. (A) Overview of the fraction of ADPr residing on histones. *n*=4 cell culture replicates, error bars represent SD. (B) Histogram visualizing the distribution of ADPr abundance across histone family members. *n*=4 cell culture replicates, error bars represent SEM. (C) Overview of the most abundant ADPr site per histone variant. (D) Multiple sequence alignment of all Histone H1 isoforms detected to be ADP-ribosylated. ADPr-modified residues are highlighted in blue. Alternative non-unique assignments of the same ADPr peptide to different histone isoforms are highlighted in red. Black and white stars indicate unique and non-unique MS detection, respectively. Black and white circles indicate presence or absence of histone ADPr via immunoblot analysis. (E) Immunoblot analysis accompanying **D**. Experiments were performed in wild-type or HPF1 KO HEK293T cells, transiently transfected with the indicated GFP-tagged histones for 24 h. H_2_O_2_ treatment was performed at 2 mM for 10 min, after which cells were lysed and GFP-IP was performed. (F) As **B**, but for Histone H1 isoforms. (G) Overview of the number of co-modified ADPr peptides (in blue) and the abundance of co-modified ADPr peptides (in grey). “ac”; acetylation, “me1”; mono-methylation, “me2”; di-methylation, “me3”; tri-methylation, “ph”; phosphorylation, “2x”; doubly-modified (including ADPr), “3x”; triply-modified (including ADPr). (H) Visualization of the fractional abundance of ADPr co-modifications occurring on Histones H1.2 and H1.4. *n*=4 cell culture replicates, error bars represent SEM

### Multiple histone variants are co-modified by ADPr and other PTMs

ADP-ribosylation can occur in the proximity of other post-translational modifications, such as phosphorylation (Hendriks et al., 2019; Tanigawa et al., 1983), as well as methylation and acetylation in the context of histones (Bartlett et al., 2018; Kassner et al., 2013; Malik and Smulson, 1984; Wong and Smulson, 1984), which could be indicative of crosstalk between the different PTMs. We investigated our data for ADPr peptides co-modified by phosphorylation, methylation, or acetylation, and in total we were able to identify 240 unique co-modified peptides corresponding to 146 co-modified sites (Fig. 4G and Table S4). Approximately 2/3^rd^ of the peptides contained two PTMs, with the other 1/3^rd^ containing three PTMs. The majority of co-modification with methylation and acetylation was found to reside on histones, whereas co-modification with phosphorylation occurred predominantly on non-histone proteins (Fig. 4G). Crosstalk around Histone H3 S11-ADPr was previously described (Bartlett et al., 2018), which correlated well with our findings in the context of HPF1 and ARH3 knockout (Fig. S4C), with acetylation of K15 overall being the major co-modification, followed by mono- and di-methylation of K10. We were able to visualize ADPr on Histone H1 to an unprecedented degree, allowing us to examine crosstalk with other PTMs. Histone H1.2 and H1.4 were the main targets of co-modification (Fig. 4H and Table S4), with S27-ADPr and S188-ADPr flanked by methylation and acetylation for H1.4 and H1.2, respectively. Strikingly, S150-ADPr was co-modified for both H1.2 and H1.4, overall representing the highest degree of crosstalk, with frequent threonine phosphorylation at −4 or simultaneously at −4 and +4. Taken together, we demonstrate serine ADP-ribosylation targets a plethora of histone variants, particularly in the absence of ARH3, and we elucidate a high degree of co-modification of histones with serine ADPr and other PTMs.

## DISCUSSION

Employing our Af1521 enrichment strategy for studying the global effects of HPF1 and ARH3 on the human ADP-ribosylome, we corroborate that ADPr is strongly induced upon depletion of ARH3 and conversely abrogated upon depletion of HPF1 (Bonfiglio et al., 2017; Fontana et al., 2017; Gibbs-Seymour et al., 2016). While HPF1 and ARH3 inversely regulate the serine ADP-ribosylome, the functional modularity is highly homogenous which contrasts previous observations related to ADPr of glutamate and aspartate (Zhen et al., 2017).

Intriguingly, while we find serine ADPr to be the major acceptor site across investigated cells, no cellular switch in target residues from serine to glutamate or aspartate was detectable upon depletion of HFP1, although this was previously shown *in vitro* (Palazzo et al., 2018). We confirmed our proteomics data by orthogonal *in vivo* approaches, demonstrating that histone variants are predominantly modified with serine ADPr (Fig. 4E, Fig. S3B, Fig. S4B). Likewise, treating cellular extracts with hydroxylamine, which chemically cleaves ADPr localized to glutamates and aspartates, did not result in discernable differences in ADPr signal across investigated cell conditions (Fig. S2C).

Our observations likely reflect differences in experimental strategies for studying PARP enzymes. For example, it may be impractical to capture *in vivo* conditions in biochemical assays considering the existence of numerous PARP1 substrates and range of modulators (Ariumi et al., 1999; Boehler et al., 2011; Chen et al., 2018; Kauppinen et al., 2006; Kun et al., 2004; Menissier de Murcia et al., 2003; Ray Chaudhuri et al., 2016; Schreiber et al., 2002). Moreover, proteomic strategies used for mapping ADPr on glutamate and aspartate are based upon derivatization of ADPr into a chemical mark (+15.011 Da) using hydroxylamine (Zhang et al., 2013). However, use of hydroxylamine can induce chemical artifacts (+15.011 Da) on acidic residues (Geiszler et al., 2020), mimicking the mark used for ADPr identification. Indeed, treatment of bovine serum albumin with hydroxylamine under native conditions revealed the same aberrant chemical marks (+15.011 Da), which could lead to erroneous identification of ADPr-modified aspartate and glutamate residues (Supplementary Note). Contrary to this, our Af1521 methodology enriches peptides covalently modified with entire ADP-ribose moieties and is therefore not prone to similar chemical artifacts.

The enrichment of ADPr-modified peptides without downstream PARG treatment furthermore allowed us to infer the cellular MARylation versus PARylation equilibrium, hereby uncovering that MARylation is a global and dominant phenomenon in human cells. While observations capture the state of the cell upon lysis, and considering that ADPr is a very dynamic modification (Polo and Jackson, 2011), the detected MARylation probably stems from substrates initially being modified with PAR, which is then rapidly degraded *in vivo* into MAR by high hydrolase activity of eraser enzymes (Bonfiglio et al., 2020; Fontana et al., 2017; Hanzlikova et al., 2020), and PARP1 itself (Rudolph et al., 2020). Still, considering this dynamic regulation of ADPr has remained unnoticed for decades, along with the importance of better understanding the endogenous ADPr equilibrium, underscores the necessity for investigating ADPr dynamics under native conditions using unbiased proteomics strategies. Collectively, this work provides ap owerful resource for researchers working on PARP1, ADPr and PARP inhibitors, and the HPF1- and ARH3-dependent ADP-ribosylome bolsters the global integrative view of cellular regulation.

## ACKNOWLEDGEMENTS

The work carried out in this study was in part supported by the Novo Nordisk Foundation Center for Protein Research, the Novo Nordisk Foundation (grant agreement numbers NNF14CC0001 and NNF13OC0006477), Danish Council of Independent Research (grant agreement numbers 4002-00051, 4183-00322A, 8020-00220B and 0135-00096B), and The Danish Cancer Society (grant agreement R146-A9159-16-S2). The proteomics technology applied were part of a project that has received funding from the European Union’s Horizon 2020 research and innovation program under grant agreement EPIC-XS-823839. We would like to thank members of the NNF-CPR Mass Spectrometry Platform for instrument support and technical assistance. Work in I.A.’s laboratory was supported by Wellcome Trust (101794, 210634); Biotechnology and Biological Sciences Research Council (BB/R007195/1); and Cancer Research United Kingdom (C35050/A22284).

## AUTHOR CONTRIBUTIONS

S.C.B-L., I.A.H., and A.K.L.F.S.R. prepared MS samples, I.A.H. and S.C.B-L. measured all samples on the mass spectrometer, processed all MS raw data, and performed data analysis. S.C.B-L. and E.P. performed IB analysis. M.L.N. conceived the project. M.L.N. and I.A. supervised the project. M.L.N., I.A.H. and S.C.B-L. wrote the manuscript with input from all authors.

## SUPPLEMENTAL FIGURE LEGENDS

**Figure S1.**
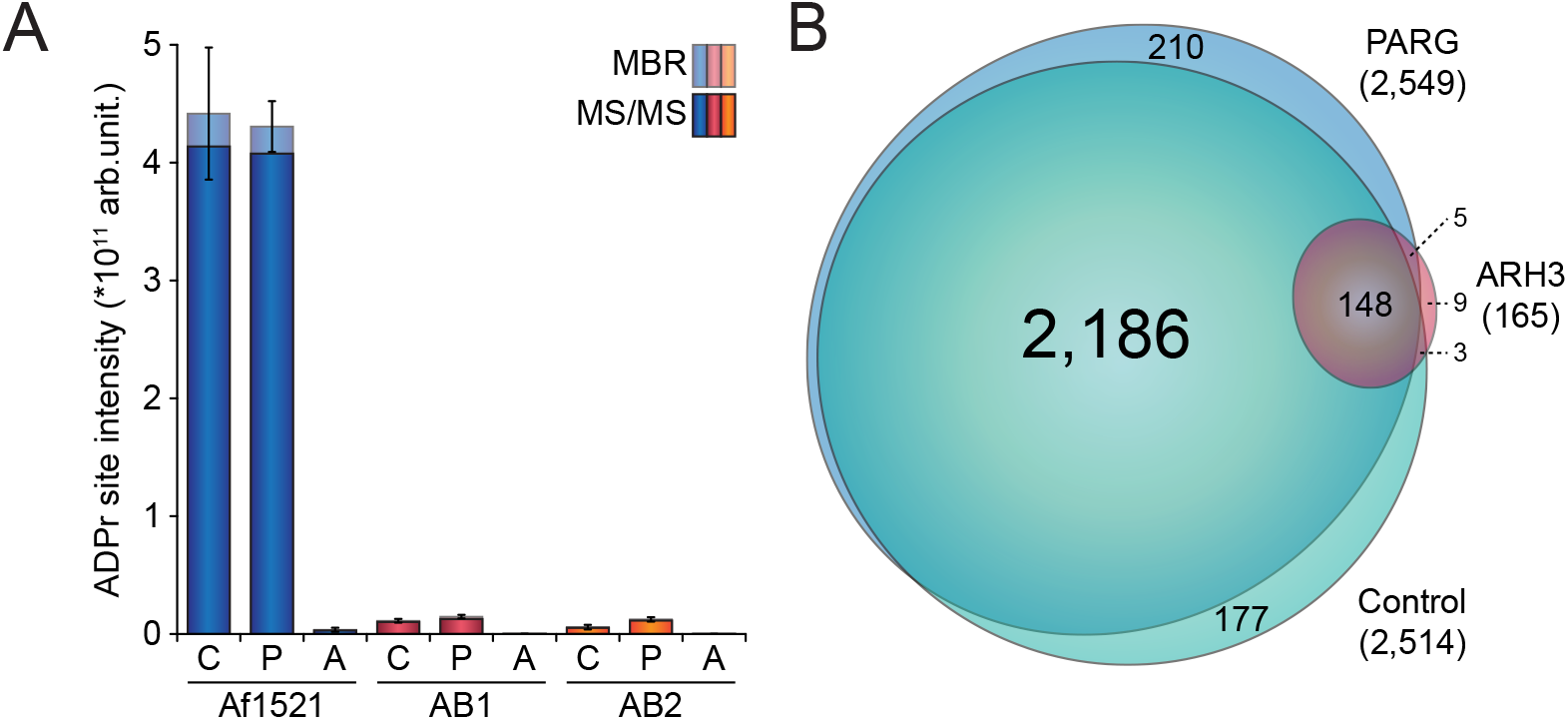
Evaluation of different erasers and enrichers for the purification of ADPr. Related to Figure 1. (A) Overview of the ADPr site abundance across experimental conditions. Error bars represent SD, *n*=4 purification replicates. “C”; control, “P”; PARG, “A”; ARH3, “MBR”; matching between runs. (B) Scaled Venn diagram depicting overlap between ADPr sites in Af1521-enriched samples.

**Figure S2.**
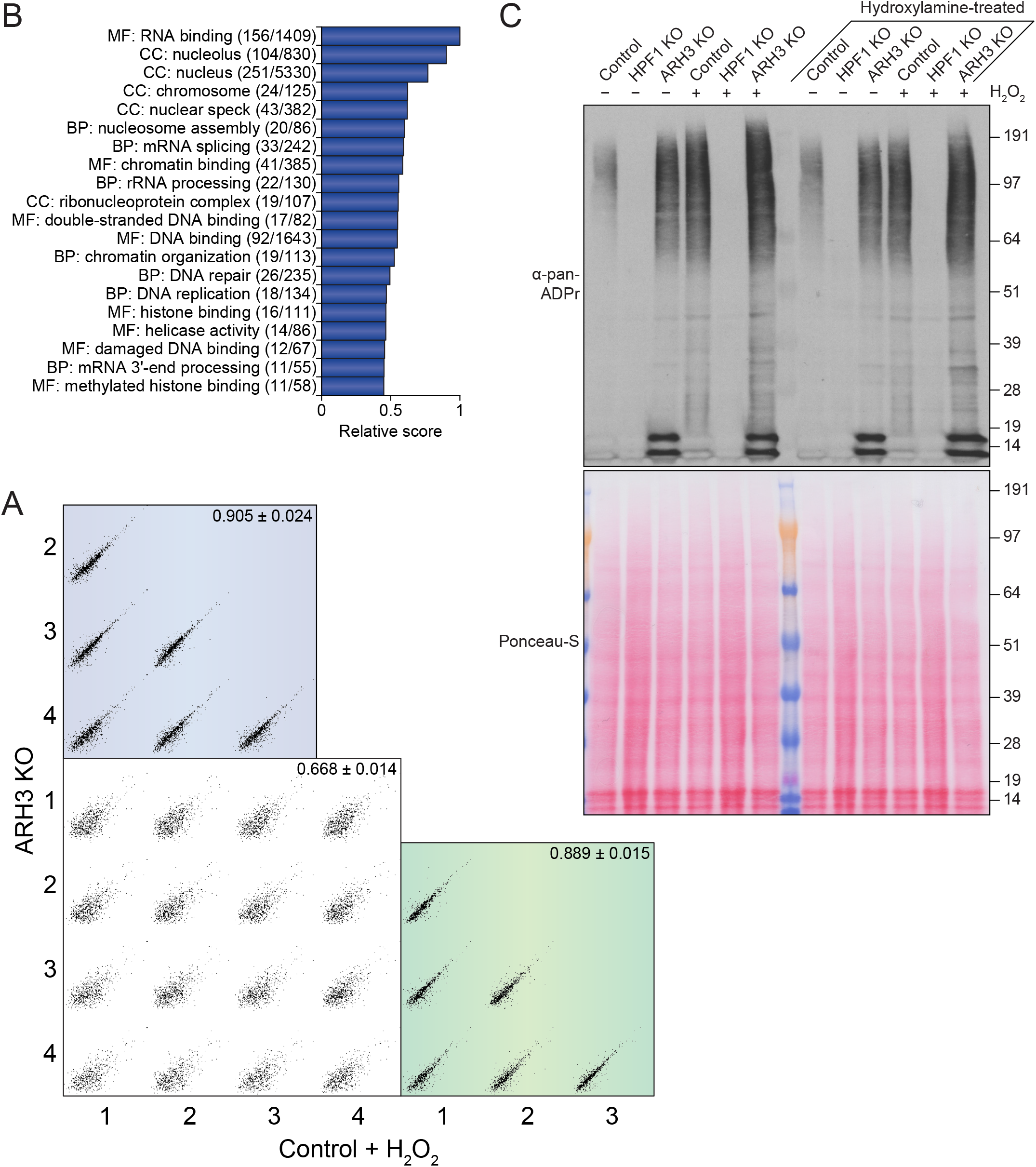
The ADP-ribosylome in HPF1 and ARH3 knockout cells. Related to Figures 2 and (A) ‘B-2 bomber plot’ visualizing the correlation of ADPr site intensities between all H_2_O_2_-treated control samples and untreated ARH3 KO samples. Numbers indicate Pearson correlation ± SD. (B) Term enrichment analysis using Gene Ontology annotations, comparing ADPr target proteins detected in both H_2_O_2_-treated control cells and untreated ARH3 KO cells, to the total proteome. Relative score is based on multiplication of logarithms derived from the enrichment ratio and the q-value. Terms were significant with q<0.02, as determined through Fisher Exact Testing with Benjamini-Hochberg correction. “BP”; biological process, “CC”; cellular compartment, “MF”; molecular function. (C) Immunoblot analysis showing that hydroxylamine treatment of lysates does not affect the distribution of ADPr. Ponceau-S analysis serves as loading control.

**Figure S3.**
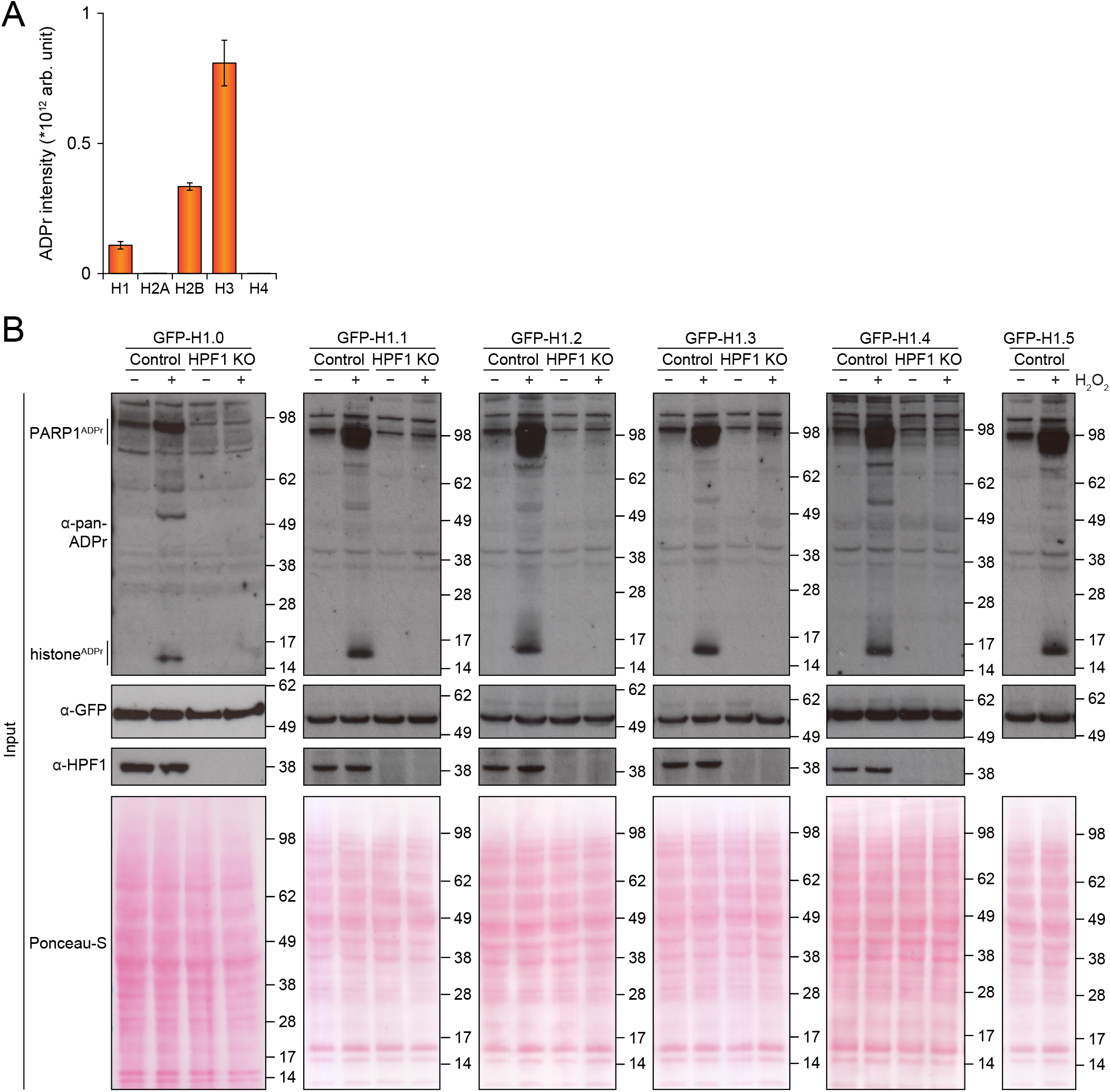
ADPr modification of histone family members. Related to Figure 4. (A) Overview of the fraction of ADPr residing on histones, merged for all experimental conditions. *n*=4 cell culture replicates, error bars represent SD. (B) Input and loading control immunoblots and Ponceau-S analysis copondinrresg to Figure 4D.

**Figure S4.**
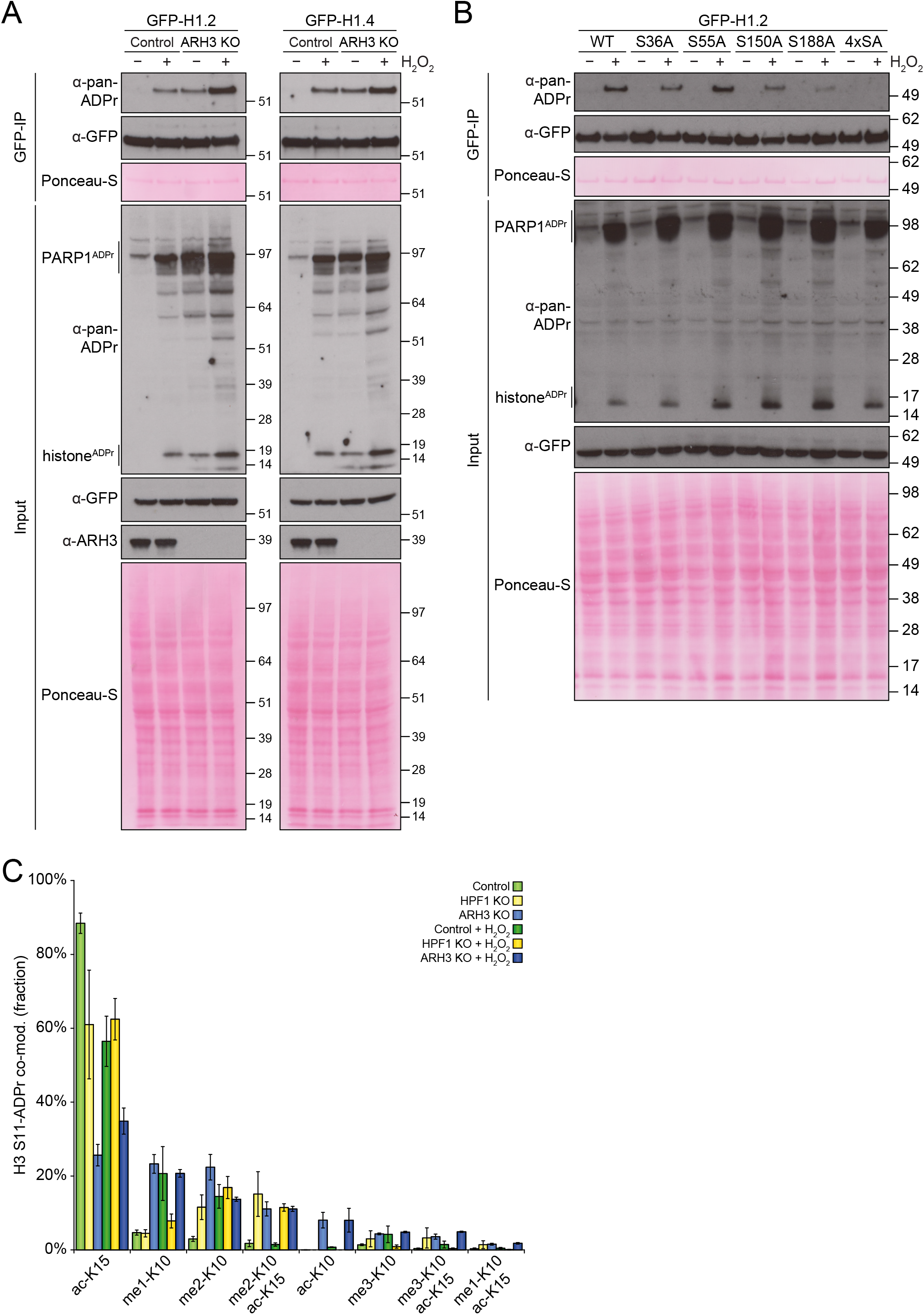
Validation of Histone H1.2 and H1.4 serine ADPr. Related to Figure 4. (A) Immunoblot experiments performed in wild-type or ARH3 KO HEK293T cells, transiently transfected with the indicated GFP-tagged histones for 24 h. H_2_O_2_ treatment was performed at 2 mM for 10 min, after which cells were lysed and GFP-IP was performed. Ponceau-S analysis serves as a loading control. (B) As **A**, but analyzing serine-to-alanine substitution mutants of GFP-tagged Histone H1.2, using wild-type HEK293T cells. “4xSA”; mutant with all four serine-to-alanine mutations. (C) Visualization of the fractional abundance of S11-ADPr co-modifications occurring on Histone H3. *n*=4 cell culture replicates, error bars represent SEM.

## STAR METHODS

### RESOURCE AVAILABILITY

#### Lead Contact

Further information and requests for resources and reagents should be directed to and will be fulfilled by the Lead Contact, Michael Lund Nielsen.

#### Materials Availability

All unique reagents generated in this study are available from the lead contact upon reasonable request.

#### Data and Code Availability

The mass spectrometry proteomics data have been deposited to the ProteomeXchange Consortium via the PRIDE (Perez-Riverol et al., 2019) partner repository with the dataset identifier PXD023835.

### EXPERIMENTAL MODEL AND SUBJECT DETAILS

#### Cell lines

HeLa cells (CCL-2, female), U-2 OS (U2OS) cells (HTB-96, female), and HEK293T cells (CRL-3216, female), were acquired via the American Type Culture Collection, and cultured at 37 °C and 5% CO2 in Dulbecco’s Modified Eagle’s Medium (Invitrogen) supplemented with 10% fetal bovine serum and a penicillin/streptomycin mixture (100 U/mL; Gibco). U2OS cells with HPF1 knockout KO (Gibbs-Seymour et al., 2016), U2OS cells with ARH3 KO (Fontana et al., 2017), HEK293T cells with HPF1 KO (Gibbs-Seymour et al., 2016), and HEK293T cells with ARH3 KO (Hanzlikova et al., 2020), were described previously. Cells were routinely tested for mycoplasma. Cells were not routinely authenticated.

#### Bacteria

BL21(DE3) Competent *E*.*coli* were acquired from New England BioLabs. Details regarding culture conditions were reported previously (Larsen et al., 2018).

#### METHOD DETAILS

#### Cell treatment

For Af1521 and pan-ADPr antibody comparison experiments, ADP-ribosylation was induced in HeLa by treatment of the cells with 1 mM H_2_O_2_ (Sigma Aldrich) for 10 min in PBS at 37°C. One batch of cells was prepared, totaling 2.7 billion cells, corresponding to approximately 75 million HeLa cells per technical replicate measured on the mass spectrometer (MS). For HPF1 KO and ARH3 KO experiments, U2OS cells were mock treated or ADP-ribosylation was induced by treatment of the cells with 1 mM H_2_O_2_ for 10 min in PBS at 37°C. For MS analysis, approximately 150 million U2OS cells were cultured per biological replicate. For immunoblot analyses, HEK293T cells were mock treated or ADPr was induced with 2 mM H_2_O_2_ in DPBS with calcium and magnesium (Gibco) for 10 min. All immunoblot analyses were performed in duplicate.

#### Cell lysis and protein digestion

The full procedure for enrichment of ADPr from cells using the Af1521 macrodomain was done as described previously (Hendriks et al., 2019; Larsen et al., 2018). Briefly, cells were washed twice with ice-cold PBS, and gently scraped at 4 °C in a minimal volume of PBS. Cells were pelleted by centrifugation at 500*g*, and lysed in 10 pellet volumes of Lysis Buffer (6 M guanidine-HCl, 50 mM TRIS, pH 8.5). Complete lysis was achieved by alternating vigorous shaking with vigorous vortexing, for 30 seconds, prior to snap freezing of the lysates using liquid nitrogen. Frozen lysates were stored at −80 °C until further processing. Lysates were thawed and sonicated at 30 W, for 1 second per 1 mL of lysate, spread across 3 separate pulses. Tris(2-carboxyethyl)phosphine and chloroacetamide were added to a final concentration of 10 mM, and the lysate was incubated for 1 hour at 30 °C. Proteins were digested using Lysyl Endopeptidase (Lys-C, 1:100 w/w; Wako Chemicals) for 3 hours, and diluted with three volumes of 50 mM ammonium bicarbonate. Samples was further digested overnight using modified sequencing grade Trypsin (1:100 w/w; Sigma Aldrich). Digested samples were acidified by addition of trifluoroacetic acid (TFA) to a final concentration of 0.5% (v/v), cleared by centrifugation, and purified using reversed-phase C18 cartridges (SepPak Classic, 360 mg sorbent, Waters) according to the manufacturer’s instructions. Elution of peptides was performed with 30% ACN in 0.1% TFA, peptides were frozen overnight at −80 °C, and afterwards lyophilized for 96 h.

#### Purification of ADP-ribosylated peptides

Lyophilized peptides were dissolved in AP buffer (50 mM TRIS pH 8.0, 1 mM MgCl_2_, 250 μM DTT, and 50 mM NaCl), after which either no enzyme, PARG enzyme (kind gift from Prof. Dr. Michael O. Hottiger), or ARH3 enzyme (kind gift from Prof. Dr. Bernhard Lüscher) were added in a 1:10,000 (w/w) ratio, overnight and at room temperature. For Af1521 and pan-ADPr antibody comparison experiments, no enzyme, PARG, or ARH3, were used as indicated in the figure legend. For HPF1/ARH3 KO experiments, PARG was added, theoretically in order to reduce ADPr polymers to monomers. GST-tagged Af1521 macrodomain was produced in-house using BL21(DE3) bacteria, and coupled to glutathione Sepharose 4B beads (Sigma-Aldrich), essentially as described previously (Hendriks et al., 2019; Larsen et al., 2018). Pan-ADPr antibodies were a kind gift from Cell Signaling Technology (CST), and 10x 100 μL batches of E6F6A (AB1) and D9P7Z (AB2) were received pre-conjugated to agarose beads. For Af1521 and pan-ADPr antibody comparison experiments, 50 μL of sepharose beads with GST-tagged Af1521 or 50 μL of antibody beads were added to the samples, and mixed at 4 °C for 3 h. For HPF1/ARH3 KO experiments, 150 μL of Af1521 beads were used. Beads were sequentially washed four times with ice-cold IP Buffer, two times with ice-cold PBS, and two times with ice-cold MQ water. On the first wash, beads were transferred to 1.5 mL LoBind tubes (Eppendorf), and LoBind tubes were exclusively used from this point on to minimize loss of peptide. Additional tube changes were performed every second washing step to minimize carryover of background. ADP-ribosylated peptides were removed from the beads using two elution steps with two bead volumes ice-cold 0.15% TFA, and the pooled elutions were cleared through 0.45 μm spin filters (Ultrafree-MC, Millipore) and subsequently through pre-washed 100 kDa cut-off filters (Vivacon 500, Sartorius). The filtered ADP-ribosylated peptides were purified using C18 StageTips at high pH (Hendriks et al., 2019), and eluted as four or five fractions for comparison and HPF1/ARH3 KO experiments, respectively. Briefly, samples were basified by adding ammonium solution to a final concentration of 20 mM, and loaded onto StageTips carrying four layers of C18 disc material (punch-outs from 47mm C18 3M™ extraction discs, Empore). Elution was sequentially performed with 4% / 10% / 25% of ACN in 20 mM ammonium hydroxide for initial experiments, and 2% / 4% / 7% / 25% of ACN in 20 mM ammonium hydroxide for HPF1/ARH3 KO experiments. The last fraction (F0) was prepared by performing StageTip purification at low pH on the flowthrough fraction from sample loading at high pH. All samples were completely dried using a SpeedVac at 60 °C, and dissolved in a small volume of 0.1% formic acid. Final samples were frozen at −20 °C until measurement.

#### Mass spectrometric analysis

All samples were measured using an Orbitrap Fusion™ Lumos™ Tribrid™ mass spectrometer (Thermo), and analyzed on 20-cm long analytical columns with an internal diameter of 75 μm, packed in-house using ReproSil-Pur 120 C18-AQ 1.9 µm beads (Dr. Maisch). On-line reversed-phase liquid chromatography to separate peptides was performed using an EASY-nLC™ 1200 system (Thermo), and the analytical column was heated to 40°C using a column oven. Peptides were eluted from the column using a gradient of Buffer A (0.1% formic acid) and Buffer B (80% ACN in 0.1% formic acid). The primary gradient ranged from 3% buffer B to 24% buffer B over 50 minutes, followed by an increase to 40% buffer B over 12 minutes to ensure elution of all peptides, followed by a washing block of 18 minutes. Electrospray ionization (ESI) was achieved using a Nanospray Flex Ion Source (Thermo). Spray voltage was set to 2 kV, capillary temperature to 275°C, and RF level to 30%. Full scans were performed at a resolution of 60,000, with a scan range of 300 to 1,750 m/z, a maximum injection time of 60 ms, and an automatic gain control (AGC) target of 600,000 charges. Precursors were isolated with a width of 1.3 m/z, with an AGC target of 200,000 charges, and precursor fragmentation was accomplished using electron transfer disassociation with supplemental higher-collisional disassociation (EThcD) at 20 NCE, using calibrated charge-dependent ETD parameters. Calibration of charge-dependent ETD parameters was essentially performed as described previously (Rose et al., 2015). Charge-dependent ETD calibration resulted in ETD activation times of 48.39 ms for z=3 precursors, 27.22 ms for z=4 precursors, and 17.42 ms for z=5 precursors. This equates to an ETD Time Constant (τ) of 2.42. Precursors with charge state 3-5 were isolated for MS/MS analysis, and prioritized from charge 3 (highest) to charge 5 (lowest). Selected precursors were excluded from repeated sequencing by setting a dynamic exclusion of 60 seconds. MS/MS spectra were measured in the Orbitrap, with a loop count setting of 5, a maximum precursor injection time of 120 ms, and a scan resolution of 60,000.

#### Data analysis

Analysis of the mass spectrometry raw data was performed using MaxQuant software (REFs), version 1.5.3.30. MaxQuant default settings were used, with exceptions outlined below. Two separate computational searches were performed, one for Af1521 and pan-ADPr antibody comparison data (“Search 1”), and the other for the HPF1/ARH3 KO data (“Search 2”). For generation of the theoretical spectral library, a HUMAN.fasta database was downloaded from UniProt on the 24^th^ of May, 2019. N-terminal acetylation, methionine oxidation, cysteine carbamidomethylation, and ADP-ribosylation on all amino acid residues known to potentially be modified (C, D, E, H, K, R, S, T, and Y), were included as variable modifications. For Search 2, additionally lysine acetylation, lysine mono-, di-, and tri-methylation, and serine, threonine, and tyrosine phosphorylation, were included as variable modifications. For the Search 2 first search, which is only used for mass recalibration, phosphorylation was omitted as a variable modification. Up to 6 or 5 missed cleavages were allowed, a maximum allowance of 4 or 3 variable modifications per peptide was used, and maximum peptide mass was set to 4,600 or 3,900 Da, respectively for Search 1 and 2. Second peptide search was enabled (default). Matching between runs was enabled with an alignment time window of 20 minutes and a match time window of 42 or 60 s, first search precursor mass tolerance of 20 or 15 ppm, main search precursor mass tolerance of 4.5 or 3 ppm, respectively for Search 1 and 2. For fragment ion masses, a tolerance of 20 ppm was used. Modified peptides were filtered to have an Andromeda score of >40 (default), and a delta score of >20. Data was automatically filtered by posterior error probability to achieve a false discovery rate of <1% (default), at the peptide-spectrum match, the protein assignment, and the site-specific levels.

#### Data filtering

Beyond automatic filtering and FDR control as applied by MaxQuant, the data was manually filtered in order to ensure proper identification and localization of ADP-ribose. PSMs modified by more than one ADP-ribose were omitted. PSMs corresponding to unique peptides were only used for ADP-ribosylation site assignment if localization probability was >0.90, with localization of >0.75 accepted only for purposes of intensity assignment of further evidences. Erroneous MaxQuant intensity assignments were manually corrected in the sites table, and based on localized PSMs only (>0.90 best-case, >0.75 for further evidences). For ADP-ribosylation target proteins table derived from the HPF1/ARH3 KO data, the proteinGroups.txt file generated by MaxQuant was filtered to only contain those proteins with at least one ADP-ribosylation site detected and localized post-filtering as outlined above, with cumulative ADP-ribosylation site intensities based only on localized evidences.

#### Plasmids and site-directed mutagenesis

The mammalian expression H1.0 GFP-tagged pDEST47 (Invitrogen) construct was generated using LR Clonase II enzyme mix (Invitrogen) from H1.0 pDONR221 vector, acquired from DNASU Plasmid Repository. H1.1, H1.2, H1.3, H1.4, H1.5 GFP-tagged constructs were kind gifts from Gyula Timinszky. H1.2 GFP-tagged point mutants were made using QuikChange Lightning Site-Directed Mutagenesis Kit (Agilent) following the manufacturer’s protocol.

#### Transfection and immunoprecipitation

HEK293T cells were plated in 10-cm dishes and transiently transfected with an indicated plasmid for 24 h using Polyfect (Qiagen) following the manufacturer’s instructions. The cells were washed with PBS and were lysed with Triton-X100 lysis buffer (50 mM Tris-HCl pH 8.0, 100 mM NaCl, 1% Triton X-100) supplemented with 5 mM MgCl_2_, protease and phosphatase inhibitors (Roche), 2 μM Olaparib, 1 μM PARGi PDD00017273 at 4°C. Protein concentrations were normalized using Bradford Protein Assay (Bio-Rad), and cell lysates were incubated with GFP-Trap MA magnetic agarose beads (ChromoTek) for 1 h while rotating at 4°C. Beads were washed five times with Triton X-100 lysis buffer and eluted with 2x NuPAGE LDS sample buffer (Invitrogen) with TCEP (Sigma).

#### Immunoblot analysis

U2OS cells were lysed in STBS buffer (2% SDS, 150 mM NaCl, 50 mM TRIS-HCl pH 8.5) at room temperature. U2OS lysates were homogenized by shaking at 99°C at 1,400 RPM for 30 min. HEK293T cells were lysed with Triton-X100 lysis buffer (50 mM Tris-HCl pH 8.0, 100 mM NaCl, 1% Triton X-100) supplemented with 5 mM MgCl_2_, protease and phosphatase inhibitors (Roche), 2 μM Olaparib, 1 μM PARGi PDD00017273 at 4°C. HEK293T lysates were incubated with 0.1% Benzonase (Sigma) for 30 min at 4°C, centrifuged at 14,000 rpm for 15 min, and the supernatants were collected. Protein concentrations were analyzed by Bradford Protein Assay (Bio-Rad). Prior to loading, lysates were supplemented with 1x NuPAGE LDS sample buffer (Invitrogen) with DTT or TCEP, and size-separated on 4-12% Bis-Tris gels using MOPS running buffer. For U2OS samples, proteins were transferred to Amersham™ Protran® nitrocellulose membranes (GE Healthcare). For HEK293T samples, proteins were transferred onto nitrocellulose membranes (Bio-Rad) using Trans-Blot Turbo Transfer System (Bio-Rad). Equal total protein loading was ensured by Ponceau-S staining. Membranes were blocked using 5% BSA solution (for U2OS samples) or 5% non-fat dried milk (for HEK293T samples) in PBS supplemented with 0.1% Tween-20 (PBST) for 1 h. Subsequently, membranes were incubated with primary antibodies overnight at 4°C, and afterwards washed three times with PBST. U2OS experiment membranes were incubated with Goat-anti-rabbit HRP conjugated secondary antibody (Jackson Immunoresearch, 111-036-045), at a concentration of 1:10,000 for 1 h at room temperature. HEK293T experiment membranes were incubated with peroxidase-conjugated secondary anti-rabbit antibody (Agilent, P0399), at a concentration of 1:3,000 for 1 h at room temperature. Membranes were washed three times with PBST prior to detection using Novex ECL Chemiluminescent Substrate Reagent Kit (Invitrogen). The following primary antibodies were used for immunoblot analysis in this study, and diluted at 1:1,000 unless stated otherwise. For U2OS experiments: Poly/Mono-ADP Ribose (E6F6A) Rabbit mAb #83732 (CST), PARP1 (46D11) Rabbit mAb #9532 (CST), HPF1 Rabbit pAb HPA043467 (Atlas Antibodies), ADPRHL2 (ARH3) Rabbit pAb HPA027104 (Atlas Antibodies), GAPDH Rabbit pAb ab9485 (Abcam). For HEK293T experiments: Pan-ADPr (MABE1016) Rabbit (Millipore); at 1:1,500, ADPRHL2 (ARH3) Rabbit pAb HPA027104 (Atlas Antibodies); at 1:2,000, GFP (ab290) Rabbit (Abcam); at 1:5,000, custom-made HPF1 antibody was previously described (Gibbs-Seymour et al., 2016).

## QUANTIFICATION AND STATISTICAL ANALYSIS

Details regarding the statistical analysis can be found in the respective figure legends. Statistical handling of the data was primarily performed using the freely available Perseus software (Tyanova et al., 2016), and includes term enrichment analysis through FDR-controlled Fisher Exact testing, density estimation for highly populated scatter plots, volcano plot analysis, hierarchical clustering, and principle component analysis. Protein Gene Ontology annotations and UniProt keywords used for term enrichment analysis were concomitantly downloaded from UniProt with the HUMAN.fasta file used for searching the RAW data. Sequence context analysis was performed using iceLogo software, version 2.1 (Colaert et al., 2009). Multiple sequence alignment was performed using Clustal Omega as integrated in UniProt, using the (default) transition matrix Gonnet, gap opening penalty of 6 bits, gap extension of 1 bit, and using the HHalign algorithm and its default settings as the core alignment engine (Söding, 2005).

## SUPPLEMENTAL ITEM TITLES

**Supplementary Table 1. Overview of ADPr sites, related to Figure 1**. A list of all 2,758 unique human ADPr sites identified in the Af1521 and pan-ADPr antibody comparison experiments, complete with qualitative and quantitative information. *n*=4 cell culture replicates.

**Supplementary Table 2. Overview of ADPr sites, related to Figures 2, 3, and 4**. A list of all 1,596 unique human ADPr sites identified in the HPF1 KO and ARH3 KO experiments, complete with qualitative and quantitative information. *n*=4 cell culture replicates.

**Supplementary Table 3. Overview of ADPr target proteins, related to Figures 2, 3, and 4**. A list of all 799 unique human ADPr target proteins identified in the HPF1 KO and ARH3 KO experiments, complete with qualitative and quantitative information. *n*=4 cell culture replicates.

**Supplementary Table 4. Overview of co-modified ADPr peptides and sites, related to Figure**

A list of all 146 unique human ADPr-co-modified sites identified in the HPF1 KO and ARH3 KO experiments, complete with qualitative and quantitative information. *n*=4 cell culture replicates.

