## Supplementary note for "The regulatory landscape of the human HPF1- and ARH3-dependent ADP-ribosylome"

Proteomics strategies used for mapping ADPr on glutamate and aspartate are based upon derivatization of ADPr into a chemical mark (+15.0109 Da) using hydroxylamine (HA) (Zhang et al., 2013). Recently, an investigation of TMT data – where HA is used as part of the sample preparation workflow – revealed the second most common unexpected mass shift to be +15.0109 Da (Geiszler et al., 2020). This mass shift could be explained via conversion of carboxylic acid to hydroxamic acid, resulting in addition of an NH group on glutamate and aspartate residues (Geiszler et al., 2020).

We wondered whether the usage of HA during ADPr samples preparation could also result in derivatization of glutamate and aspartate residues towards hydroxamic acid, and to which extent this would occur. To this end, we prepared a technical standard using commercial Bovine Serum Albumin (BSA), a routinely used quality control protein for mass spectrometric analysis, and a protein which is highly unlikely to be substantially ADP-ribosylated. We subjected BSA to an ADPr sample processing workflow essentially as described previously (Li et al., 2019), while including or excluding the HA treatment step (Supplementary Note Figure A). The resulting BSA peptides were analyzed using an Orbitrap Exploris™ 480 mass spectrometer using higher-energy collisional dissociation (HCD) fragmentation.

Overall, in the absence of HA treatment, we did not observe any hydroxamic acid modification on BSA (Supplementary Note Figure B-E). Strikingly, when BSA was treated with HA, we observed dozens of hydroxamic acid modifications on aspartate and glutamate residues, overall accounting for 8% of all identified spectra and 4% of total BSA abundance. Concomitantly, we also observed a large induction of asparagine and glutamine oxidation after HA treatment (Supplementary Note Figure F-I), which was not observed for untreated BSA. Methionine oxidation, which is a common side effect during electrospray ionization (Morand et al., 1993), was unaltered between untreated and HA-treated BSA, showing an absence of technical bias. Our sequence coverage of BSA was ~50%, and we found that nearly half of all profiled aspartate and glutamate residues were modified by hydroxamic acid when BSA was treated with HA (Supplementary Note Figure J).

In summary, we demonstrate that HA treatment of BSA, a protein unrelated to ADPr regulation, results in readily detectable amounts of hydroxamic acid (+15.0109 Da) modifications on aspartate and glutamate residues. In the context of ADPr proteomics, we would like to raise awareness of this phenomenon with the community, so that future ADPr proteomics experiments can be more carefully controlled to avoid potential artificial misidentification of modified aspartate and glutamate residues.

### FIGURE LEGEND

(A) Overview of experimental design. (B) Visualization of the number of peptide-spectrum-matches (PSMs) identified with hydroxamic acid on aspartate and glutamate residues. *n*=4 technical replicates. Error bars represent SEM. (C) As **B**, but showing the fraction of modified PSMs in relation to all PSMs. (D) Visualization of the abundance (intensity, arb. unit) of peptides modified with hydroxamic acid. (E) As **D**, but showing the fraction of modified abundance in relation to total abundance. (F) Visualization of the number of PSMs identified with oxidation of asparagine, glutamine, and methionine residues. (G) As **F**, but showing the fraction of modified PSMs in relation to all PSMs. (H) Visualization of the abundance (intensity, arb. unit) of peptides modified with oxidation. (I) As **H**, but showing the fraction of modified abundance in relation to total abundance. (J) Schematic representation of BSA, visualizing which part of the sequence was detected (in blue). Aspartate and glutamate residues which were modified by hydroxamic acid after HA treatment are indicated in red, non-modified residues are indicated in grey.

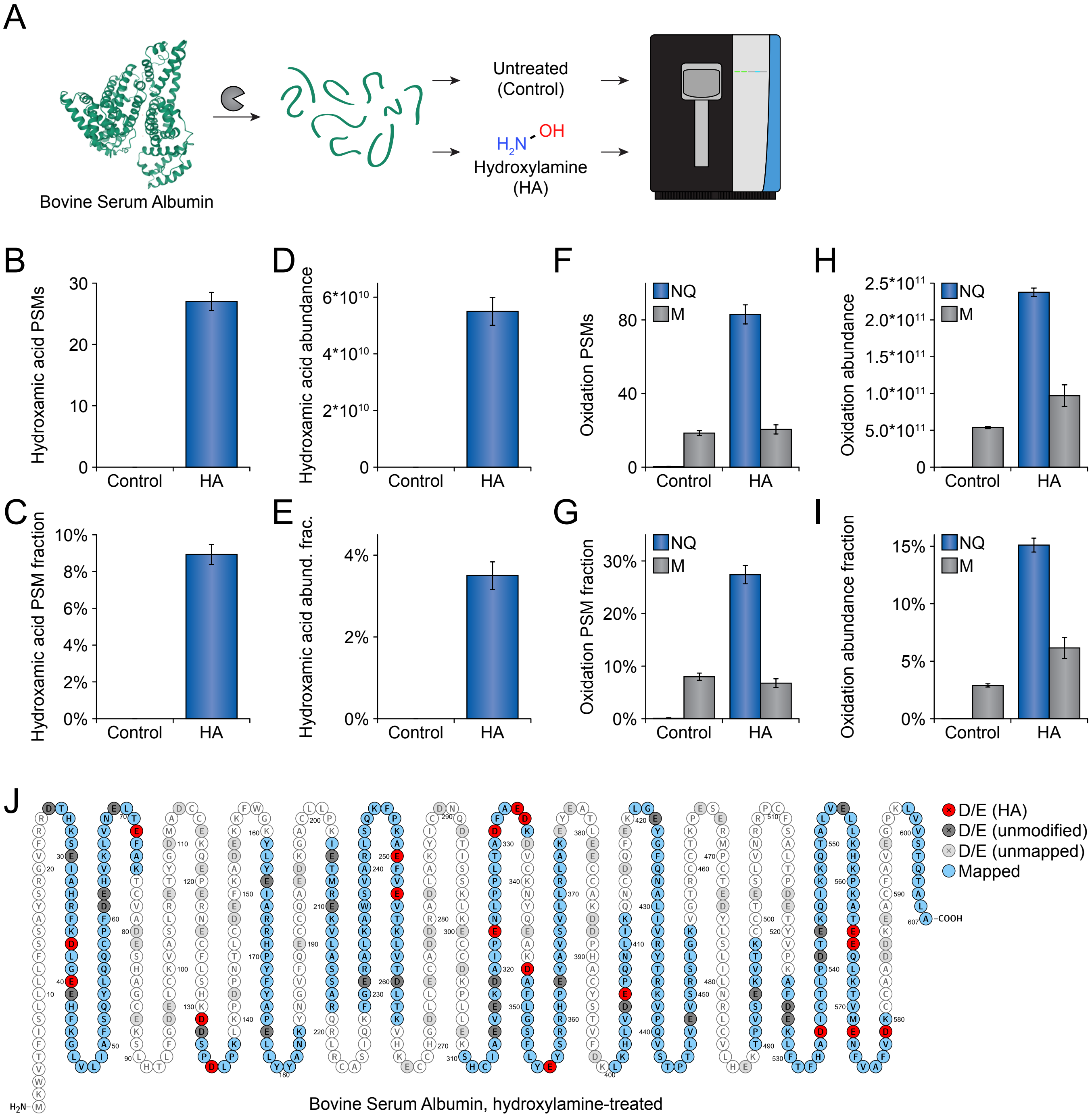

A
Bovine Serum Albumin
Untreated
(Control)
OH
$H_2N$
Hydroxylamine
(HA)
B
Hydroxamic acid PSMs
30
20
10
0
Control
HA
C
Hydroxamic acid PSM fraction
10%
8%
6%
4%
2%
0%
Control
HA
D
Hydroxamic acid abundance
6*10^10
4*10^10
2*10^10
0
Control
HA
E
Hydroxamic acid abund. frac.
4%
2%
0%
Control
HA
F
Oxidation PSMs
NQ
M
80
40
0
Control
HA
G
Oxidation PSM fraction
NQ
M
30%
20%
10%
0%
Control
HA
H
Oxidation abundance
NQ
M
2.5*10^11
2.0*10^11
1.5*10^11
1.0*10^11
5.0*10^11
0
Control
HA
I
Oxidation abundance fraction
NQ
M
15%
10%
5%
0%
Control
HA
J
D/E (HA)
D/E (unmodified)
D/E (unmapped)
Mapped
$H_2N$–
–COOH
Bovine Serum Albumin, hydroxylamine-treated
